# A fast and accurate algorithm to test for binary phenotypes and its application to PheWAS

**DOI:** 10.1101/109876

**Authors:** Rounak Dey, Ellen M. Schmidt, Goncalo R. Abecasis, Seunggeun Lee

**Affiliations:** Department of Biostatistics, University of Michigan, Ann Arbor, MI 48109, USA; Center for Statistical Genetics, University of Michigan, Ann Arbor, MI 48109, USA

## Abstract

The availability of electronic health record (EHR)-based phenotypes allows for genome-wide association analyses in thousands of traits, and has great potential to identify novel genetic variants associated with clinical phenotypes. We can interpret the phenome-wide association study (PheWAS) result for a single genetic variant by observing its association across a landscape of phenotypes. Since PheWAS can test 1000s of binary phenotypes, and most of them have unbalanced (case:control = 1:10) or often extremely unbalanced (case:control = 1:600) case-control ratios, existing methods cannot provide an accurate and scalable way to test for associations. Here we propose a computationally fast score test-based method that estimates the distribution of the test statistic using the saddlepoint approximation. Our method is much faster than the state of the art Firth’s test (∼ 100 times). It can also adjust for covariates and control type I error rates even when the case-control ratio is extremely unbalanced. Through application to PheWAS data from the Michigan Genomics Initiative, we show that the proposed method can control type I error rates while replicating previously known association signals even for traits with a very small number of cases and a large number of controls.

## Introduction

Over the last decade, genome wide association studies (GWASs) have proved instrumental to unravelling the genetic complexities of hundreds of diseases and traits and their associations with common genomic variations. To date, thousands of GWASs have identified more than 4000 significant loci to be associated with human diseases and traits.^1^ However, since most GWASs investigate a single disease or trait, they cannot exploit the cross-phenotype associations or pleiotropy^2^ where a single genetic variant can be associated with multiple phenotypes. Phenome-wide association study (PheWAS) has been proposed as an alternative approach to take advantage of the pleiotropy phenomenon by studying the impact of genetic variations across a broad spectrum of human phenotypes or ‘phenome’. It is a complementary approach to GWAS in the sense that while GWAS attempts to identify phenotype-to-genotype associations, PheWAS uses a genotype-to-phenotype approach. The first PheWAS^3^ was published as a proof-of-principle study, which demonstrated that the PheWAS strategy could be applied to successfully identify the expected gene-disease associations. Additional studies^4-8^ have shown that the PheWAS approach can further identify novel disease-SNP associations.^9^

The PheWAS approach depends on the availability of detailed phenotypic information. Currently, most of the PheWASs are applied to clinical cohorts linked to electronic health records (EHR) and utilize the International Classification of Disease (ICD) billing codes to define clinical phenotypes. The ICD codes provide an intuitive ordering of the phenotypes based on clinical disease and trait classifications. Since the current genotyping and imputation technologies allow for genotyping tens of millions of variants at very low cost,10 an extensive PheWAS can attempt to investigate the genotype-phenotype associations by performing genome-wide association analyses in thousands of traits. We can interpret the PheWAS result of a single genetic variant by observing its associations across the phenome. Such a PheWAS is exhaustive in nature and has great potential to identify novel variants associated with clinical diseases.

One of the main challenges of the PheWAS analysis is that most of the phenotypes are binary phenotypes with unbalanced (1:5) or often extremely unbalanced (1:600) case-control ratios (See *Figure S1)*, since the data is collected in cohorts. Although standard asymptotic tests, such as the Wald, score and likelihood ratio tests, are relatively well calibrated and asymptotically equivalent^11^for common variants (minor allele frequency (MAF) > 0.05) in balanced case-control studies, they can inflate type I error for low frequency (0.01 < MAF ≤ 0.05) and rare variants (MAF ≤ 0.01) in unbalanced case-control studies.12 Moreover, since the Wald and likelihood ratio tests need to calculate the likelihood or the maximum likelihood estimator under the full model, which is computationally expensive, they are not scalable for the amount of tests that PheWASs attempt. On the other hand, the score test is computationally efficient as it does not need to calculate the maximum likelihood under the full model. However, as mentioned before, it suffers from having highly inflated type I error rates in unbalanced studies. Ma *et al*. proposed Firth’s penalized likelihood ratio test13 as a solution to control the type I error rates in such situations. Firth’s test, despite being well calibrated and robust for testing low frequency and rare variants in unbalanced studies, lacks in computational efficiency as it also involves calculating the maximum likelihood under the full model. For instance, the projected computation time of the Firth’s test to test 1500 phenotypes across 10 million SNPs is ∼ 117 CPU-years (2000 cases, 18000 controls). Thus, it is impractical to apply the Firth’s test for analyzing large PheWAS datasets.

In this paper, we propose a score-based single variant test for binary phenotypes which is well calibrated for controlling the type I errors and can adjust for covariates even in extremely unbalanced case-control studies. Moreover, our test is computationally efficient and scalable to test thousands of phenotypes across millions of SNPs in large PheWAS datasets. Our proposed test (SPA) is based on the score statistics and estimates the null distribution using the saddlepoint approximation^14-16^ instead of the normal approximation17 traditionally used in score tests. We further develop an improvement of our test (fastSPA) which renders the most computationally challenging steps to be dependant only on the number of carriers (subjects with at least one minor allele) rather than the sample size. This improved test can substantially reduce the computation time, especially for low frequency and rare variants where the number of carriers is very low compared to the sample size. The projected computation time of our method to test for 1500 phenotypes across 10 million SNPs is ∼ 400 CPU-days (2000 cases, 18000 controls) which is more than a 100 times improvement over Firth’s test. In addition, through the extensive simulation studies and analysis of the Michigan Genomics Initiative (MGI) data, we demonstrate that the proposed approach can control type I errors and is powerful enough to replicate known association signals.

## Material and Methods

### Logistic regression model and saddlepoint approximation method

We consider a case-control study with sample size *n*. For the *i*^*th*^ subject, let *Y*_*i*_ = 1 or 0 denote the case-control status, *X*_*i*_ the *k* × 1 vector of non-genetic covariates including the intercept, and *G*_*i*_ the number of minor alleles (*G*_*i*_ = 0,1,2) of the variant to test. To relate genotypes to phenotypes, we use the following logistic regression model, 
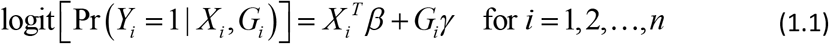
where *β* is a *k* × 1 vector of coefficients of the covariates, and *γ* is the genotype log-odds ratio. Under this model, we are interested in testing for the genetic association by testing the null hypothesis *H*_0_: *γ* = 0. Let 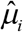 be the estimate of *μ*_*i*_ = Pr(*Y*_*i*_ = 1|*X_i_*), which is a probability to be a case under *H*_0_. A score statistic for *γ* from the model (1.1) is given by 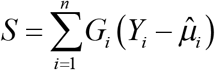. Suppose 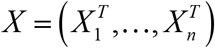 is the *n* × *k* matrix of covariates, *G* = (*G*_1_,…,*G*_*n*_)^*T*^ is the genotype vector, *W* is a diagonal matrix with the *i*^*th*^ diagonal element being 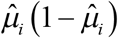, and 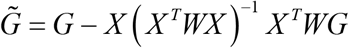 is a covariate adjusted genotype vector in which covariate effects are projected out from the genotypes (details given in the Appendix). Then *S* can be written as

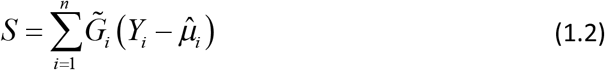
and the mean and variance of *S* under *H*_0_ are *E*_*H*_0__ (*S*) = 0 and 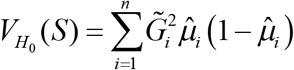, respectively, where 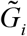 is the *i*^*th*^ element of 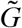.

The traditional score test approximates the null distribution using a normal distribution which depends only on the mean and the variance of the score statistic. The *p*-value can be obtained by comparing the observed test statistic, *s* and *N*(0,*V*_*H*_0__(*S*)). Normal approximation works well near the mean of the distribution, but performs very poorly at the tails. The performance is especially poor when the underlying distribution is highly skewed, such as in unbalanced case-control outcomes12, since normal approximation cannot incorporate higher moments such as skewness. In addition, the convergence rate of normal approximation^18-20^ is *O*(*n*^-1/2^), which is not fast enough for rare variants.

Saddlepoint approximation was introduced by Daniels^14^ as an improvement over the normal approximation. Contrary to normal approximation, where only the first two cumulants (mean and variance) are used to approximate the underlying distribution, saddlepoint approximation uses the entire cumulant generating function. Jensen^21^ further showed that saddlepoint approximation has a relative error bound of *O*(*n*^-3/2^) making it a considerable improvement over the normal approximation.

To use the saddlepoint approximation, we first derive the cumulant generating function of *S* from the fact that *Y*_*i*_ ∼ *Bernoulli*(*μ*_*i*_) under *H*_0_. Let 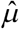 be an *n* × 1 vector with the *i*^*th*^ element being 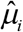 From (1.2), the estimate of the cumulant generating function of the score statistic *S* is, 
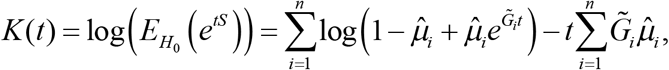
and the estimate of the first and second order derivatives of *K* are 
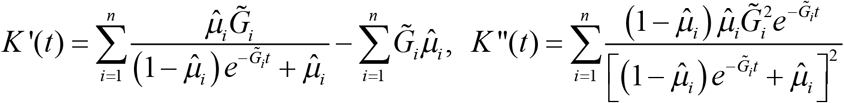
respectively. We note that *K,K*′ and *K*″ are plug-in estimates in which we plug in 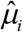 instead of *μ*_*i*_. Then, according to the saddlepoint method (Barndorff-Nielson^15; 16^), the distribution of *S* at *s* can be approximated by,

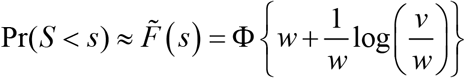
where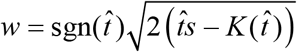, 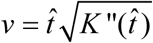 is the solution to the equation 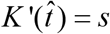, and Φ is the distribution function of a standard normal distribution.

### Implementation details and approaches to reduce the computation time

The saddlepoint approximation method involves finding the root of the saddlepoint equation *K*’(*t*) = *s*. It is easy to verify that *K*’ is strictly increasing as *K*”(*t*) > 0 for all –∞ < *t* <∞, and 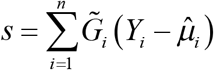 lies between 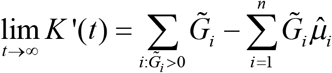 and 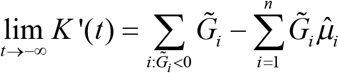.

Therefore a unique root exists, and we can use popular root-finding algorithms (Newton-Raphson,^22; 23^ bisection,^23^ secant,^23^ Brent’s method^24^) to efficiently solve this equation. For our simulation studies and real-data applications we applied a combination of the Newton-Raphson and bisection method to solve the saddlepoint equations.

The most computationally demanding step in this saddlepoint approximation method is calculating the cumulant generating function and its derivatives. Here we propose several approaches to reduce the computational complexities associated with these calculations.

#### Faster calculation of the CGF using a partially normal approximation approach

The most computationally intensive step in the saddlepoint method is the calculation of the cumulant generating function *K* and its derivatives. In each step of the root-finding algorithm we need to calculate *K*,*K*’ and *K*”, each of which needs *O*(*n*) computations. Using the fact that many elements of *G* are zeroes (i.e, homozygous major genotypes), we propose a fast computation method that speeds up the computation to *O*(*m*), where *m* is the number of non-zero elements in *G*. Without loss of generality we assume that the first *m* subjects have at least one minor allele each and rests have homozygous major genotypes. We can then express *S* as *S* = *S*_1_ + *S*_2_ where 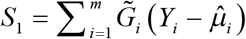 and 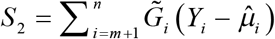. Let *Z* = (*X^T^WX*)^−1^ and *Z_l_* be the *l*^*th*^ element of *Z*. Then we can further express *S*_2_ as,

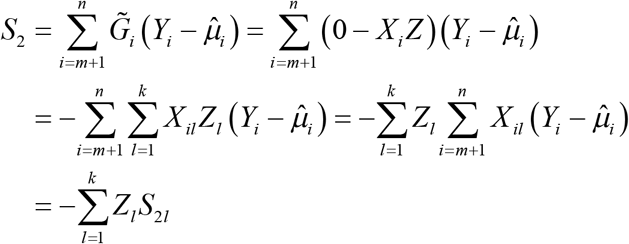
where 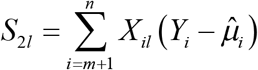. Now, if we assume that the non-genetic covariates are relatively balanced in the sample, then the normal distribution should be a good approximation for the null distribution of each *S*_2*l*_. Since *S*_2_ is a weighted sum of the *S*_2*l*_s, we can also approximate the null distribution of *S*_2_ using a normal distribution with mean and the variance under *H*_0_ given by *E*_*H*_0__ (*S*_2_) = 0 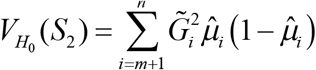. Then, the CGF of *S*_2_ can be approximated by, 
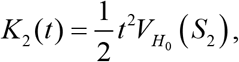
and the CGF of *S* = *S*_1_ + *S*_2_ can be approximated by, 
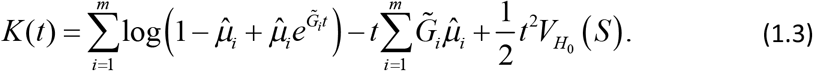

In order to calculate the first two terms at the right hand side of (1.3), we will need 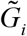 s for *i* = 1,…,*m*, which can be calculated in *O*(*m*) computations since *G* only has *m* many non-zero elements and the quantity *X* (*X*^*T*^*WX*)^−1^*X^T^W* can be pre-calculated. Then, the first two terms will require only *O*(*m*) computations as both of them sums over *m*many elements. Next, the variance *V*_*H*_0__ (*S*) can be further broken down into,

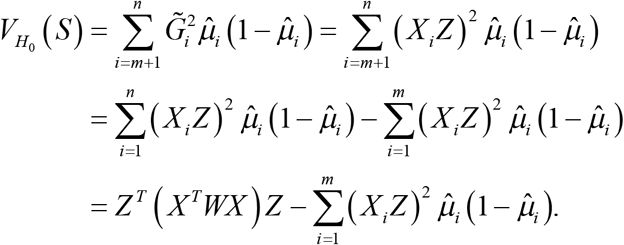

Since *X*^*T*^*WX* can be pre-calculated and *Z* is a *k* × 1 vector, the first term requires *O*(*k*) computations, and the second term requires *O*(*m*) computations, which implies that the calculation of *V*_*H*_0__ (*S*_2_) requires *O*(*m*) calculations assuming *k* < *m*, i.e, the number of non-genetic covariates is smaller than the number of subjects with at least one minor allele each. Hence, the cumulant generating function *K*(*t*) can be calculated in *O*(*m*) computations. Using similar arguments, we can further show that the derivatives *K*′(*t*) and *K*″(*t*) can also be calculated in *O*(*m*) computations. Therefore, this partially normal approximation reduces the computational complexity of our test from *O*(*n*) to *O*(*m*), which is especially useful for rare variants, where *m* is much smaller than *n*.

#### Using normal approximation near the mean for faster computation

Since the normal approximation behaves well near the mean of the distribution, we can use it to obtain the *p*-value when the observed score statistic ( *s*) lies close to the mean (zero). Moreover, saddlepoint approximation can be numerically unstable very close to the mean of the distribution. Such situations can also be avoided by using normal approximation near the mean. One possible approach is to use a fixed threshold in which we apply normal approximation to obtain the *p*-value if the absolute value of the observed score statistic, |s| < *r*σ where 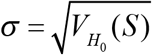 and *r* is a pre-specified value. For example, we used *r* = 2 in our simulation studies and real data analyses. For a given level *α*, this approach does not inflate type I error rates if *r* < Φ^−1^ (1 − *α* / 2), where Φ^−1^ is the inverse function of the standard normal distribution function, Φ*x*).

Alternatively, we can adaptively select the threshold using the error bound of the normal approximation given by the Berry-Esseen theorem. Suppose we are interested in controlling the type I error rate at level *α*. Let *F*_*n*_(*x*) be the true distribution function of the standardized score test statistic 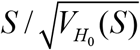. Then, according to Berry-Esseen theorem^18–20^, the maximum error bound in approximating *F*_*n*_(*x*) by Φ(*x*) is 
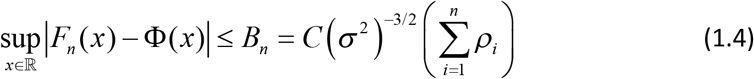
where 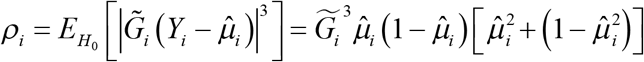, *C* is a constant. As of now, the best known estimate for *C* is 0.56, given by Shevtsova.25 Suppose *p*_*F*_ and *p*_*N*_ are *F*_*n*_(*x*) and Φ(*x*) based *p*-values. From the Berry-Esseen theorem, we can show *p*_*N*_ ≤ *P*_*F*_ + *B*_*n*_. Suppose *q* = *B*_*n*_ + *α* / 2 and *r*_*α*_ = Φ^−1^(1 — *q*). Then *p*_*N*_ ≥ *q* indicates *p*_*F*_ ≥*α*/2. Therefore, we use *r*_*α*_σ as a threshold at level *α* in which we will apply normal approximation if |s| < *r*_*α*_σ.

### Numerical Simulations

To evaluate the computation times, type I error rates and power of the proposed method, we carried out extensive simulation studies. We considered three different case-control ratios: balanced with 10000 cases and 10000 controls, moderately unbalanced with 2000 cases and 18000 controls, and extremely unbalanced with 40 cases and 19960 controls. For each choice of case-control ratios, the phenotypes were simulated based on the following logistic model, 
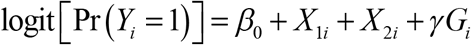
where the two non-genetic covariates *X*_1*i*_ and *X*_2*i*_ were simulated from *X*_1*i*_ ∼ *Bernoulli*(0.5) and *X*_2*i*_ ∼ *N*(0,1). The intercept *β*_0_ is chosen to correspond to prevalence 0.01. The genotype *G*_*i*_ s were generated from a *Binomial*(2, *p*) distribution where *p* was the minor allele frequency (MAF). The parameter *γ* represents the genotype log odds-ratio.

To estimate computation times and type I error rates in realistic scenarios, the MAF (p) was randomly sampled from the MAF distribution in the MGI data. For the computation time comparisons, we simulated 10^4^ variants with *γ* = 0. For the type I error comparisons, we simulated 10^9^ variants with *γ* = 0 and recorded the number of rejections at *α* = 5×10^−5^ and 5 × 10^*–8*^. We also used fixed MAFs to evaluate the effect of MAFs to computation time and type I error rates. For the power calculations, we considered two different choices for MAF, *p* = 0.01 and 0.05, and wide ranges of *γ* (*Figure 4*). For each choice of *p* and *γ* we generated 5000 variants.

**Figure 4:**
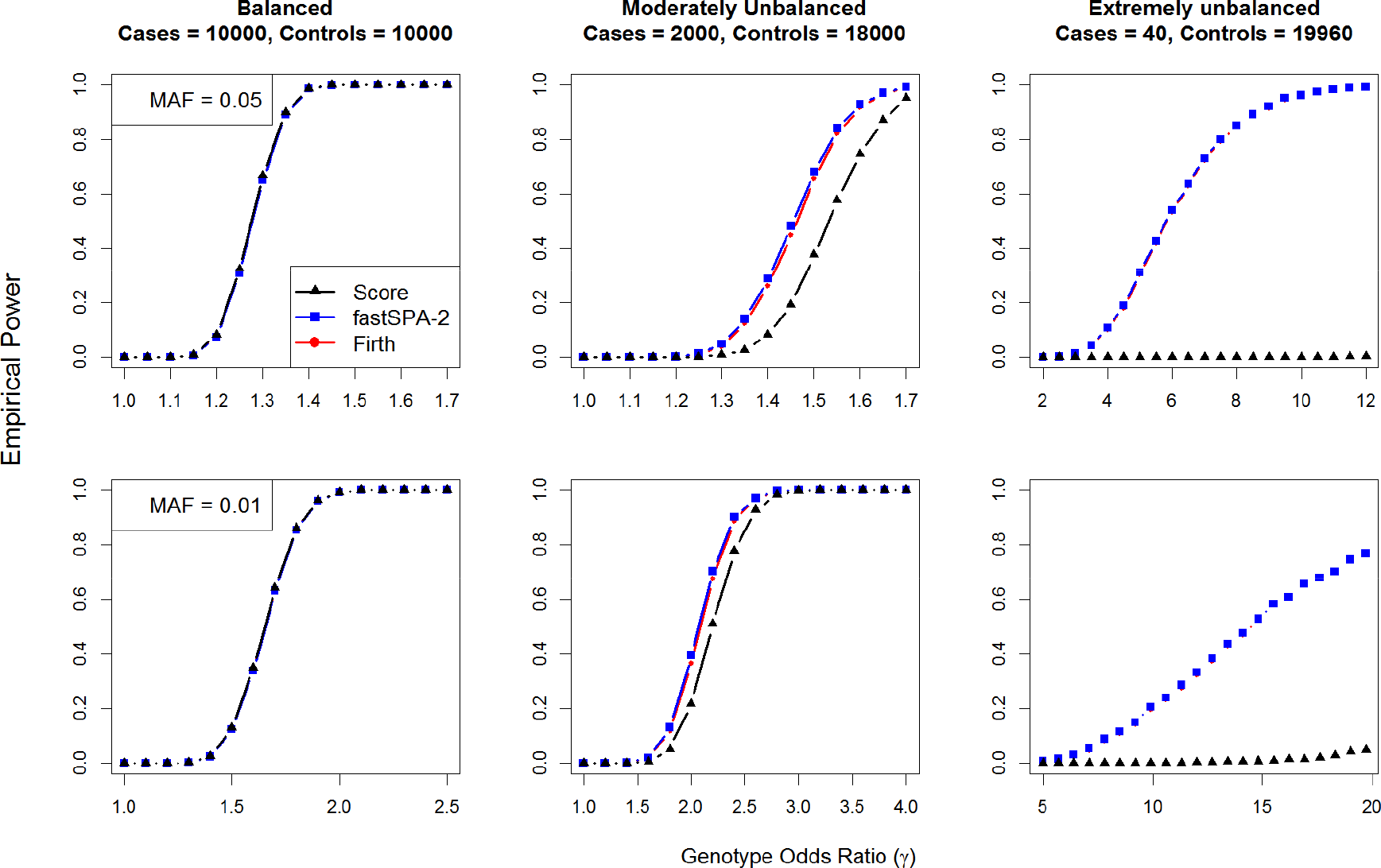
Empirical power curves for the traditional score, fastSPA-2 and Firth tests. Top panel considers MAF = 0.05 and bottom panel considers MAF = 0.01. From left to right, the plots consider case:control = 10000:10000, 2000:18000 and 40:19960, respectively. In each plot x-axis represents genotype odds ratios and y-axis represents the empirical power. Empirical power was estimated from 5000 simulated datasets at their test-specific *α* levels where their empirical type I errors are equal to 5 × 10^−8^.

We compared the computation times of seven different tests: traditional score test using normal approximation (Score); the saddlepoint approximation based test with the standard deviation threshold at 0.1 and 2 (SPA-0.1 and SPA-2); the fast saddlepoint approximation based test with the partially normal approximation improvement and the standard deviation threshold at 0.1 and 2 (fastSPA-0.1 and fastSPA-2); the fastSPA test with the Berry-Esseen bound threshold at level 5 × 10^−8^ (fastSPA-BE); and the Firth’s penalized likelihood test (Firth). Next, we compared the empirical type I errors and power curves for fastSPA-2, score and Firth tests at level 5 × 10^−8^. Since performing the Firth test 10^9^ times, which is required to estimate type I error rates at level 5 × 10^−8^, is practically impossible due to the heavy computational burden of the Firth test, we performed a hybrid approach in which we used the Firth test only when the fastSPA-2 p-values were smaller than 5 × 1 0^−3^. For the power comparison, since the score test has extremely inflated type I errors in the unbalanced and extremely unbalanced case-control scenarios (as shown in the Results section), it may not be appropriate to directly compare the power of the score test to the other two tests at the same nominal *α* level. In order to provide a more meaningful comparison, we compared their powers at their empirical *α* levels where their empirical type I errors become 5 × 10^−8^. The empirical *α* levels were selected based on the type I error simulations with variants simulated with MAF randomly sampled from the MAF distribution of the MGI data. This approach is similar to performing resampling (e.g., permutation) to control family-wise error rates. We also estimated the powers at the nominal fixed *α* = 5 × 10^−8^. In order to compare the p-values resulted from different tests, we also simulated5 ×10^6^ variants with MAFs randomly sampled from the MAF distribution of the MGI data. We further compared the inflation factors of the genomic controls at different *p*-value quantiles for fastSPA-2, fastSPA-BE and fastSPA-0.1 in order to explore the effect of the standard deviation threshold on the inflation factor.

#### Michigan Genomics Initiative (MGI) data application

To illustrate the performance of the proposed methods in real data application, we analyzed four selected phenotypes in the MGI data. The main goal of MGI is to create an institutional repository of genetic data together with rich clinical phenotypes for a broad portfolio of future medical research. DNA from blood samples of > 20,000 surgical patients at the University of Michigan Health System was genotyped (with their informed consent) on the Illumina HumanCoreExome v12.1 array, which is a combination GWAS plus exome array comprised of > 500,000 single nucleotide polymorphisms. Genotypes of the Haplotype Reference Consortium^26^ (chromosome 1-22: HRC release 1; chromosome X: HRC release 1.1) were imputed into the phased MGI genotypes (SHAPEIT2^27^ on autosomal chromosomes and Eagle2^28^ on chromosome X) using Minimac3.^29^ Excluding variants with low imputation quality (R^2^ < 0.3) resulted in dense mapping at over 39 million quality-imputed genetic markers.

Phenotypes derived from 8,940 ICD-9 billing codes were classified into 1,815 PheWAS disease states of shared disease etiology, of which 1,448 had at least 20 cases. Standard code translations were used to convert the taxonomy of diagnostic ICD-9 codes into PheWAS code groups (PheWAS code translation table version 1.2^30^). Cases were derived from electronic health records for patients with at least 2 encounters with an ICD-9 billing code. This is a typical example of many large scale PheWASs that are being conducted in recent days. In order to compare our proposed fastSPA-2 test with the traditional score test and the current gold standard Firth test in analysing such PheWAS data, we performed genome-wide association analyses for 4 selected traits, Skin Cancer (PheWAS code: 172), Type-2 diabetes (PheWAS code: 250.2, [MIM: 125853]), Primary Hypercoagulable state (PheWAS code: 286.81, [MIM: 188055]) and Cystic Fibrosis (PheWAS code: 499, [MIM: 219700]), in 18,267 unrelated individuals of European ancestry, with adjustment for age, sex, and four principal components. Genotyped samples with any missing covariate information were excluded from analysis. Since imputation quality is low for very rare variants^26^, we excluded the imputed variants with MAF < 0.001 in our main analysis, which resulted in 13 million variants. For the Firth test, we used the hybrid approach used in the type I error simulation in which the Firth test was performed only when the fastSPA-2 p-value was smaller than 5 × 10^−3^.

## Results

### Numerical Simulations

We examine the computation time, type I error control and power of the proposed fastSPA and two existing approaches, score and Firth tests, across ranges of case-control imbalance and MAFs.

#### Comparison of computation times

The projected computation times for testing 1500 phenotypes across 10 million variants using different testing methods are presented in Figure 1. To obtain computation time under realistic scenarios of the MAF distribution, the MAFs of the simulated SNPs were randomly sampled from the MAF spectrum of the MGI data (*Figure S2*). The fastSPA-2 test performs 100-300 times faster than the Firth’s test. In the unbalanced case-control setup of 2000 cases and 18000 controls, for example, the Firth’s test takes 117 CPU-years whereas fastSPA-2 only takes 1.09 CPU-years to analyze 10 million SNPs across 1500 phenotypes. This indicates that on a cluster with 100 CPU cores, the proposed test would require 4 days (without data reading) but the Firth’ test would need more than a year. When we compare fastSPA and SPA, fastSPA-0.1 performs 4-6 times faster than SPA-0.1 (e.g. 2.90 vs 12.32 CPU years when case:control = 2000:18000), and fastSPA-2 performs 1.5-2 times faster than SPA-2 (e.g. 1.09 vs 1.62 CPU years when case:control = 2000:18000). Expectedly, the computation time for fastSPA-BE is in between the computation times for fastSPA-2 and fastSPA-0.1. fastSPA-BE performs 1.3-1.8 times faster than fastSPA-0.1 and 1.6-2.8 times slower than fastSPA-2 (eg. 1.09, 1.86, 2.9 CPU years for fastSPA-2, fastSPA-BE and fastSPA-0.1 when case:control = 2000:18000).

**Figure 1:**
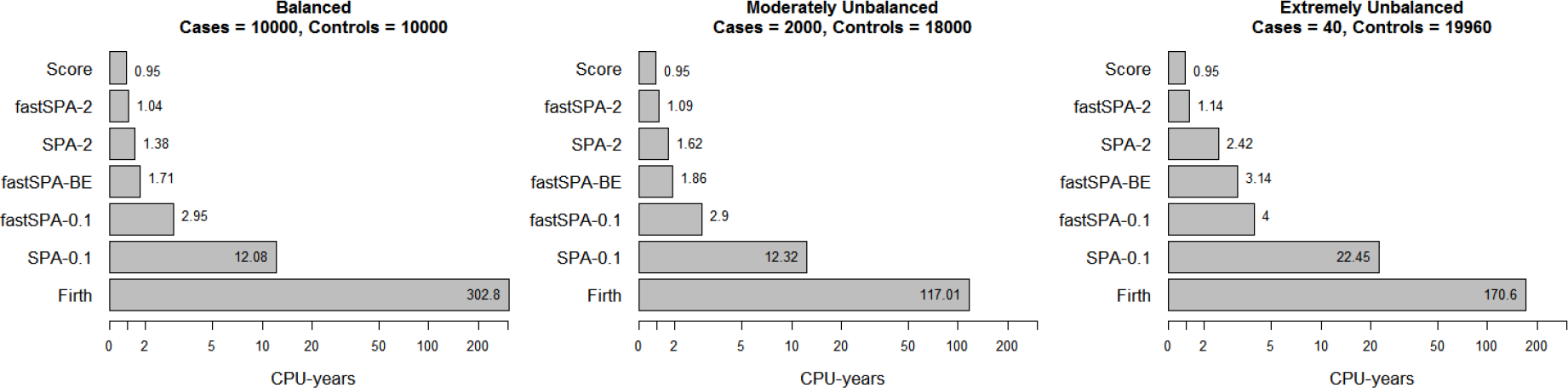
The projected computation times for testing 10 million variants across 1500 phenotypes using various tests with MAFs sampled from the MAF distribution of the MGI data. The computation times are based on testing 10000 simulated variants on an Intel i7 2.70GHz processor, and then projecting it onto a PheWAS study with 10 million variants and 1500 phenotypes.

We also recorded the computation times for variants with three different fixed MAFs 0.1, 0.01 and 0.001 in order to assess the effect of MAF on the performance of the tests. Similar to Figure 1, Table 1 also shows the superior performance of fastSPA-2 compared to all other tests. Moreover, while the computation time of SPA increases with decreasing MAFs, which may be due to the slow convergence caused by the discrete nature of the underlying distribution, fastSPA requires less computation time for rarer variants (smaller MAFs) compared to more common variants (larger MAFs). This demonstrates the potential of the partially normal approximation improvement in terms of faster computation of the *p*-values, especially for low-frequency and rare variants.

**Table 1:** Computation times for various tests when testing 10000 simulated variants with different MAFs. All computation times are in CPU-seconds on an Intel i7 2.70GHz processor.

| Case: Control | MAF | Score | SPA-0.1 | fastSPA-0.1 | fastSPA-BE | SPA-2 | fastSPA-2 | Firth |
| --- | --- | --- | --- | --- | --- | --- | --- | --- |
| 10000:10000 | 0.1 | 20 | 214 | 75 | 37 | 28 | 23 | 7251 |
|  | 0.01 | 19 | 225 | 38 | 35 | 27 | 20 | 6918 |
|  | 0.001 | 19 | 242 | 33 | 36 | 30 | 20 | 5304 |
| 2000:18000 | 0.1 | 21 | 256 | 84 | 37 | 36 | 24 | 3940 |
|  | 0.01 | 20 | 284 | 39 | 36 | 35 | 21 | 4312 |
|  | 0.001 | 19 | 326 | 34 | 41 | 40 | 20 | 3804 |
| 40:19960 | 0.1 | 21 | 376 | 98 | 70 | 38 | 24 | 3615 |
|  | 0.01 | 20 | 477 | 42 | 58 | 44 | 21 | 3598 |
|  | 0.001 | 20 | 647 | 38 | 51 | 79 | 21 | 3525 |

**Table 2:**
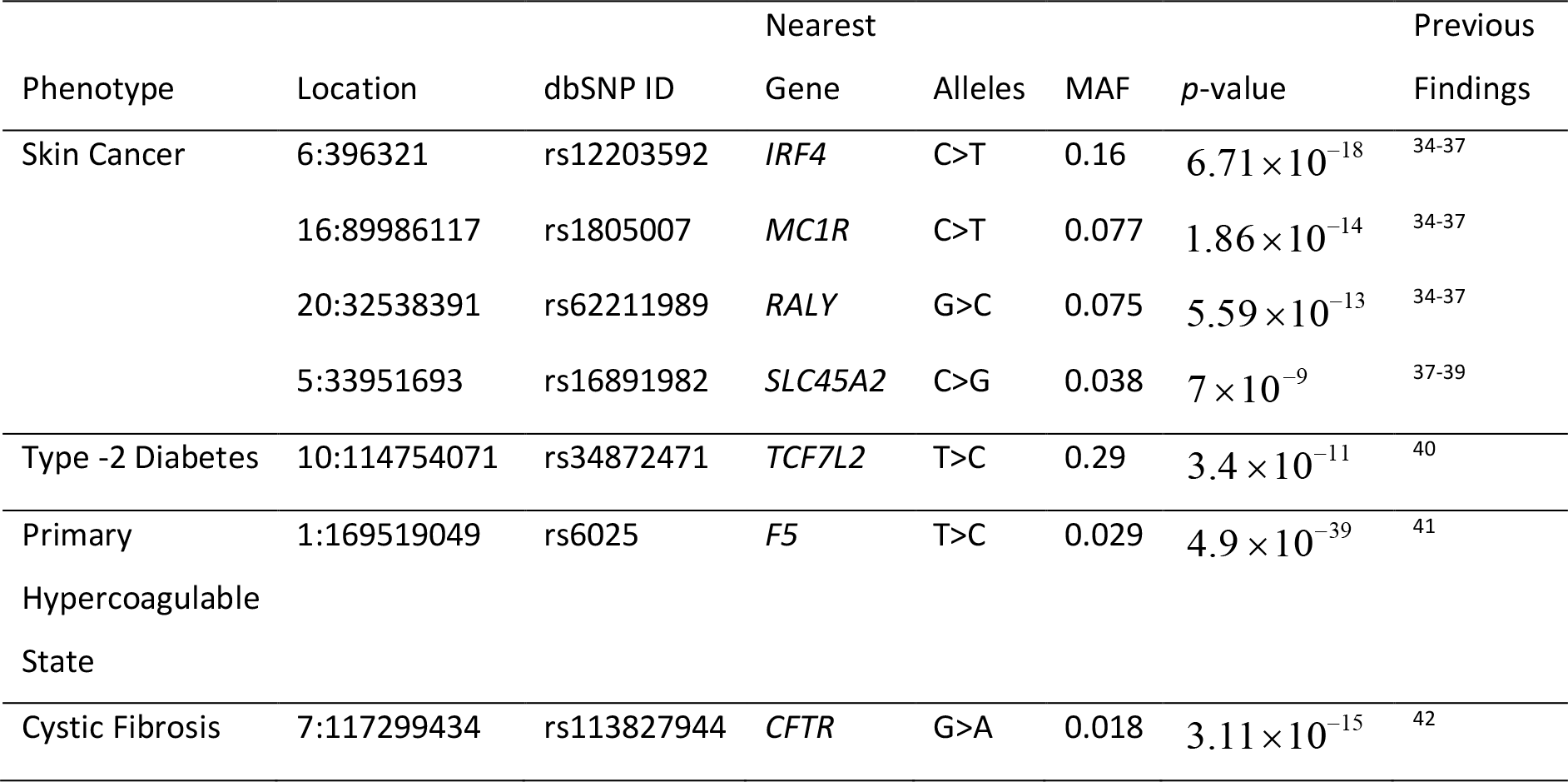
Significant SNP-phenotype associations based on fastSPA-2 test on MGI data and previous findings confirming such associations.

#### Type I error comparison

The type I error rates from 10^9^ simulated datasets are presented in Figure 2. Due to the heavy computation burden for testing these extremely large numbers of datasets, in this comparison, we only considered the traditional score test, fastSPA-2, and the hybrid version of the Firth test, in which we used the Firth test only when the fastSPA-2 p-values were smaller than 5 × 10 ^−3^. We note that both fastSPA-2 and Firth tests had well calibrated QQ plots up to 10^−6^ p-values (Figure 5), and whenever fastSPA-2 p-values > 5 × 10^−3^, Firth test p-values > 4.8 × 10^−4^ (see P-value and inflation factor comparison section), indicating that the hybrid approach can provide very accurate estimate of the type I error rates of the Firth test at very stringent α levels.

**Figure 2:**
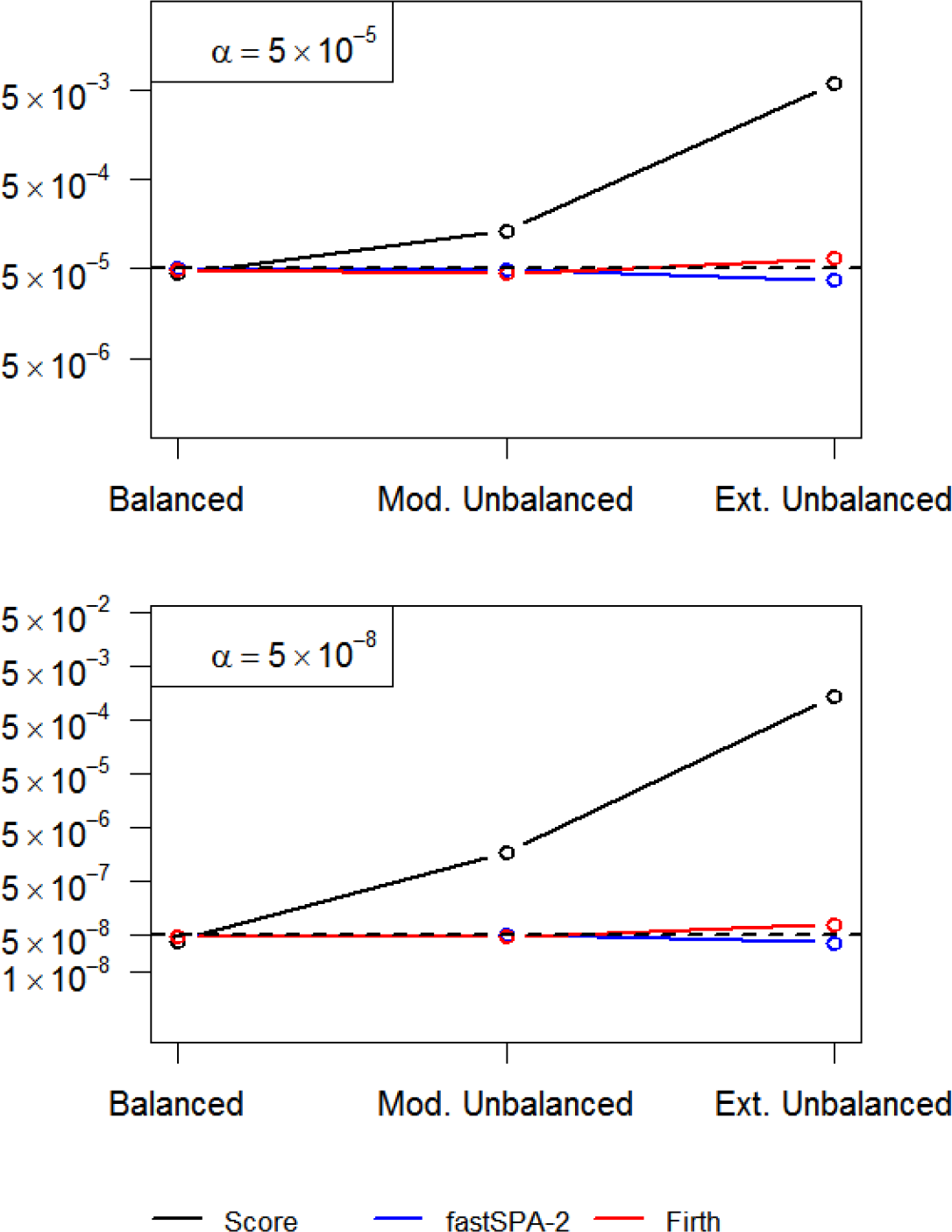
Type I error comparison between the traditional score test, fastSPA-2 and Firth tests for variants simulated with MAFs sampled from the MAF distribution of the MGI data. Type I error rates were estimated based on 10^9^ simulated datasets. From left to right on the x-axis, the plots consider case:control = 10000:10000 (Balanced), 2000:18000 (Moderately Unbalanced) and 40:19960 (Extremely Unbalanced), respectively. The top and the bottom panels show empirical type I error rates at *α* = 5 × 10^−5^ and 5 × 10^−8^ levels respectively.

**Figure 5:**
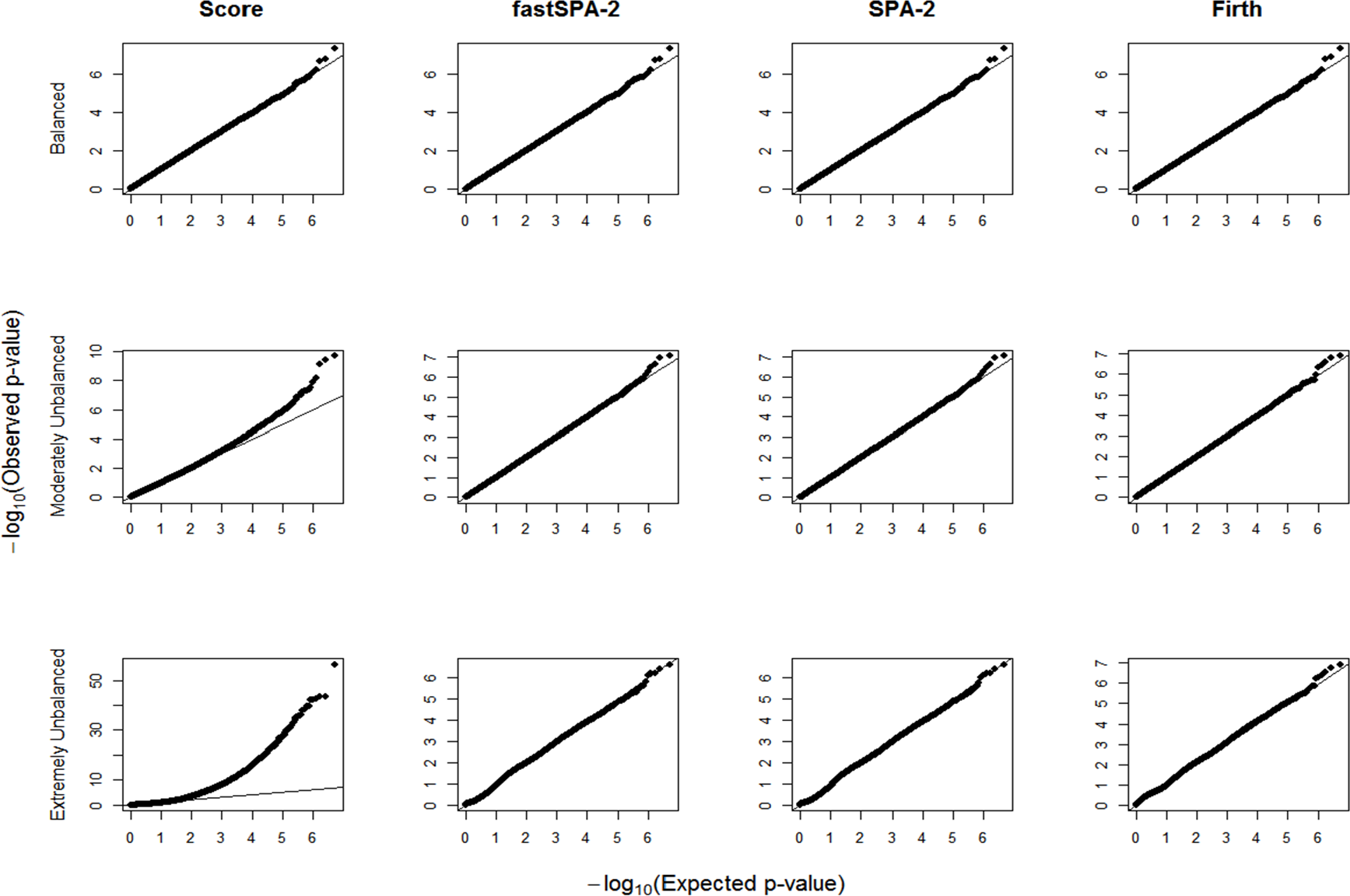
QQ plots for the traditional score, fastSPA-2, SPA-2 and Firth tests on 5 × 10^6^ simulated variants with MAF randomly sampled from the MAF distribution of the MGI data. The top, middle and bottom panels show QQ plots in the balanced (case:control = 10000:10000), moderately unbalanced (case:control = 2000:18000) and extremely unbalanced (case:control = 40:19960) case-control scenarios respectively. In each plot, x-axis represents –log10 expected p-values, and y-axis represents –log10 observed p-values.

The traditional score test had greatly inflated type I error rates for moderately unbalanced and extremely unbalanced case-control ratios, whereas fastSPA-2 can control the type I error in such situations. At the genome-wide significance level of *α* = 5 × 10^−8^, for example, the empirical type I error rates of the score test were 32 (1.63 × 10^−6^, when case:control = 2000:18000) and 26600 ( 1.33 × 10^−3^, when case:control = 40:19960) times higher than the nominal *α* = 5 × 10^−8^. In contrast, the fastSPA-2 had empirical type I error rates nearly identical (4.9 × 10^−8^, when case:control = 2000:18000) or slightly lower (3.5 × 10^−8^, when case:control = 40:19960) than the nominal *α* = 5 × 10^−8^. The Firth test also had well controlled type I error rates in the balanced and moderately unbalanced case-control scenarios ( 4.7 × 10^−8^ and 4.9 × 10^−8^, respectively at *α* = 5 × 10^−8^). Interestingly, it shows slight inflation (7.8 × 10^−8^ at *a* = 5 × 10^−8^) in the extremely unbalanced scenario. We also estimated empirical type I error rates at six different MAFs (Figure 3). The score test had deflated type I error rates for low-frequency and rare variants for the balanced case-control ratio and inflated and extremely inflated type I error rates for moderately and severely unbalanced case-control ratios. The fastSPA-2 method had overall well controlled type I error rates regardless of MAFs and case-control ratios. The Firth test had either well controlled or slightly conservative type I error rates when the case-control ratio was balanced or moderately unbalanced. However, when the case control ratio was extremely unbalanced, the Firth test had inflated type I error rates especially when the minor allele count was small (eg. *1.33 × 10*^−7^ and 1.47 × 10^−7^ for MAF = 0.0005 and 0.001 respectively at *α* = 5 × 10^∓8^ when case:control=40:19960).

**Figure 3:**
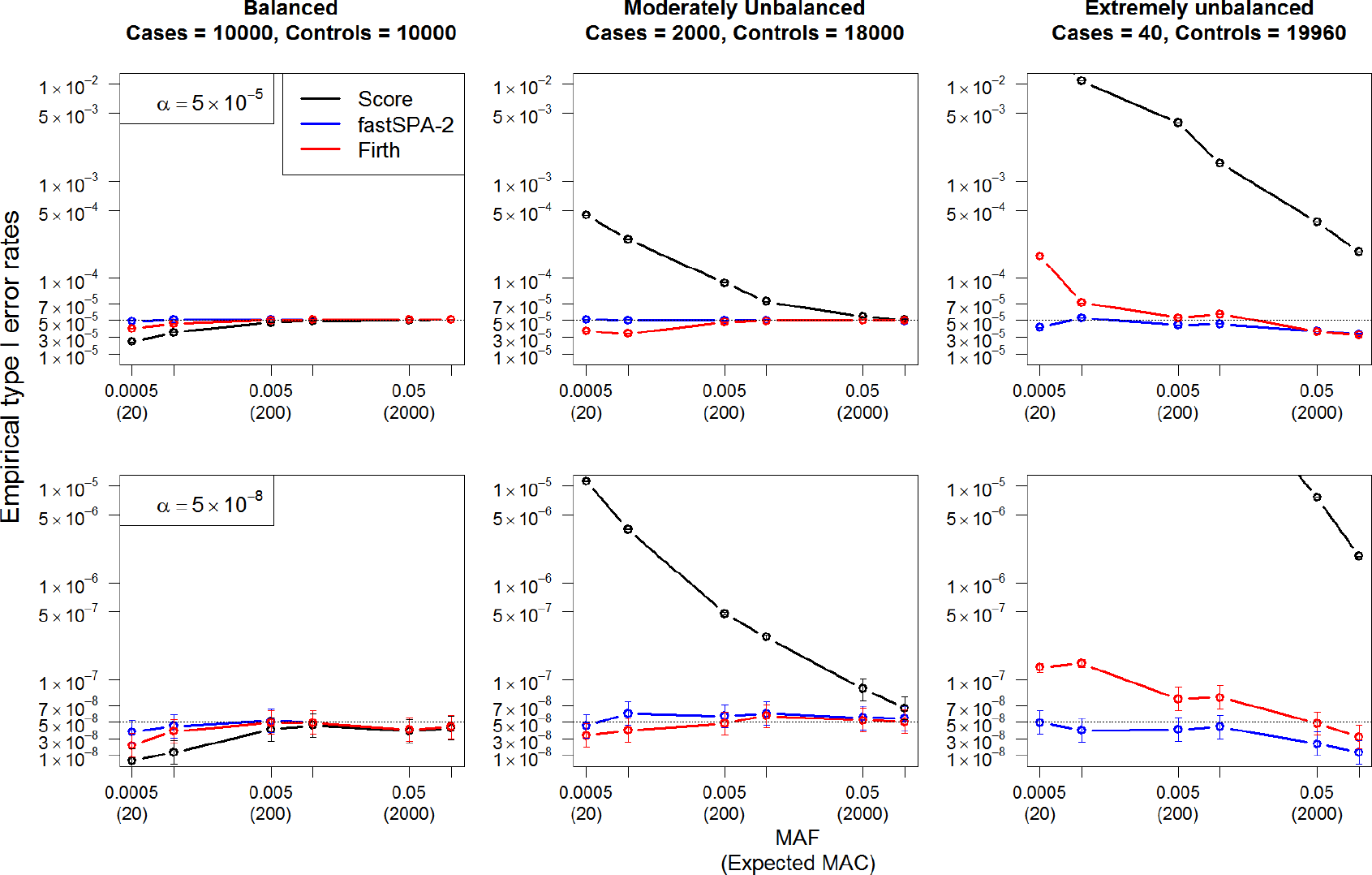
Type I error comparison at different MAFs between the traditional score test, fastSPA-2 and Firth tests. The top and bottom panels show empirical type I error rates at *α* = 5 × 10^−5^ and *α* = 5 × 10^−8^, respectively. From left to right, the plots consider case:control = 10000:10000, 2000:18000 and 40:19960, respectively. In each plot x-axis represents MAF with expected minor allele count (MAC) in the parenthesis, and y-axis represents empirical type I error rates. Empirical type I error rates were estimated based on 10^9^ simulated datasets. 95% confidence intervals at different MAFs are also presented.

#### Power comparison

Next, we compared the power curves of fastSPA-2, score and Firth tests. Note that the Firth test is a current gold standard method.^13^ Since the traditional score test had greatly inflated type I error rates, we compared the empirical powers of different tests at their test-specific empirical *α* levels. Figure 4 shows power by odds ratios when the MAF of the variant was 0.05 (top panel) and 0.01 (bottom panel). As expected, the power is higher when the case-control ratio is balanced. The empirical powers of fastSPA-2 and the Firth test were nearly identical for all case-control ratios and MAFs, which suggests that our proposed test does not suffer from any loss in power compared to the Firth test. The empirical powers of the score test were almost identical to those of fastSPA-2 and Firth test for the balanced case-control ratio. However, the score test showed substantially low power than the other two tests for the unbalanced case-control ratios due to the very small empirical *α* levels, and the power gap is especially large when the case-control ratio is extremely unbalanced. The simulation results clearly show that the proposed approach improves power over the score test when type I error rates were properly controlled. When we used nominal *α* = 5 × 10^−8^ level instead of the empirical *α* levels, score test had higher power than the other two approaches as expected (*Figure S3*), since its type I error rates were not controlled.

#### p-value and inflation factor (λ) comparison

To compare *p*-value distributions of various tests, we generated QQ plots and calculated the inflation factor (λ) of the genomic control. Figure *5* suggests strong deflation (smaller than expected) in the *p*-values based on the traditional score test in the moderately unbalanced and extremely unbalanced case-control setups, whereas fastSPA-2, SPA-2 and Firth tests resulted in well calibrated QQ plots, which suggest that these methods can control for type I errors. Moreover, the minimum Firth p-value was 4.8 × 10^−4^ for the variants with fastSPA-2 p-value > 5 × 10^−3^ among all case-control setups, which justifies our hybrid approach of performing Firth test only when fastSPA-2 p-value < 5 × 10^−3^ in the type I error simulation studies.

None of fastSPA-2, fastSPA-BE and fastSPA-0.1 tests showed any inflation or deflation in genomic controls (λ) in the balanced and moderately unbalanced case-control setups (*Table S1*). In the extremely unbalanced case-control setup, fastSPA-2 resulted in greatly deflated inflation factor (λ = 0.48) at the median of *p*-value (q = 0.5). Interestingly fastSPA-BE and fastSPA-0.1 resulted in inflated λ (both having λ = 1.83) at q = 0.5, which may be due to the discrete nature of p-values. When λ was measured at *p*-value quantiles q = 0.01 and 0.001, however, all three tests provided λ very close to unity.

### MGI Data Analysis

We applied the traditional score test, Firth test and the fastSPA-2 method to the MGI data with four phenotypes, Skin Cancer, Type-2 diabetes, Primary Hypercoagulable state, and Cystic Fibrosis, which were selected based on case-control ratios. Skin Cancer (2359 cases, 15265 controls) and Type-2 diabetes (1987 cases, 14906 controls) were moderately unbalanced, whereas Primary Hypercoagulable state (168 cases, 16401 controls) and Cystic Fibrosis (28 cases, 18212 controls) were extremely unbalanced phenotypes.

The Manhattan plots (Figure *6*) show that the traditional score test produced a large number of potentially spurious associations for all of these phenotypes, whereas all of the significant variants from our proposed test at the genome-wide significant level of *α* = 5 × 10^−8^ can be verified as truly associated with the phenotypes based on previous findings (Table *2*). In the analysis of Skin Cancer, variants in or near *IRF4* (MIM: 601900), *MC1R* (MIM: 155555), *RALY* (MIM: 614663) and *SLC45A2* (MIM: 606202) were significant at *α* = 5 × 10^−8^ and all of these four genes were previously identified as associated with pigmentation traits and skin cancers.^34–39^ In the other traits, variants in *TCF7L2* (MIM: 602228), *F5* (MIM: 612309) and *CFTR* (MIM: 602421) were significantly associated with Type2 diabetes,^40^ Primary Hypercoagulable State^41^ and Cystic Fibrosis,^42^ respectively, and all of these genes are well known to be associated with the risk of each disease. The QQ plots (Figure 7) also suggest that the *p*-values based on the traditional score test are much smaller than expected, especially for low-frequency and rare variants, whereas the *p*-values based on fastSPA-2 closely follow the uniform distribution. We also observed the Manhattan plots (*Figure S4*) including the imputed variants with MAF < 0.001 in the analysis. The inclusion of rarer variants resulted in extreme inflation in the number of potentially spurious associations for the traditional score test. However, our proposed test still produced none to very few new associations. The Manhattan plots and QQ plots for the Firth test were almost identical to those of our proposed test.

**Figure 6:**
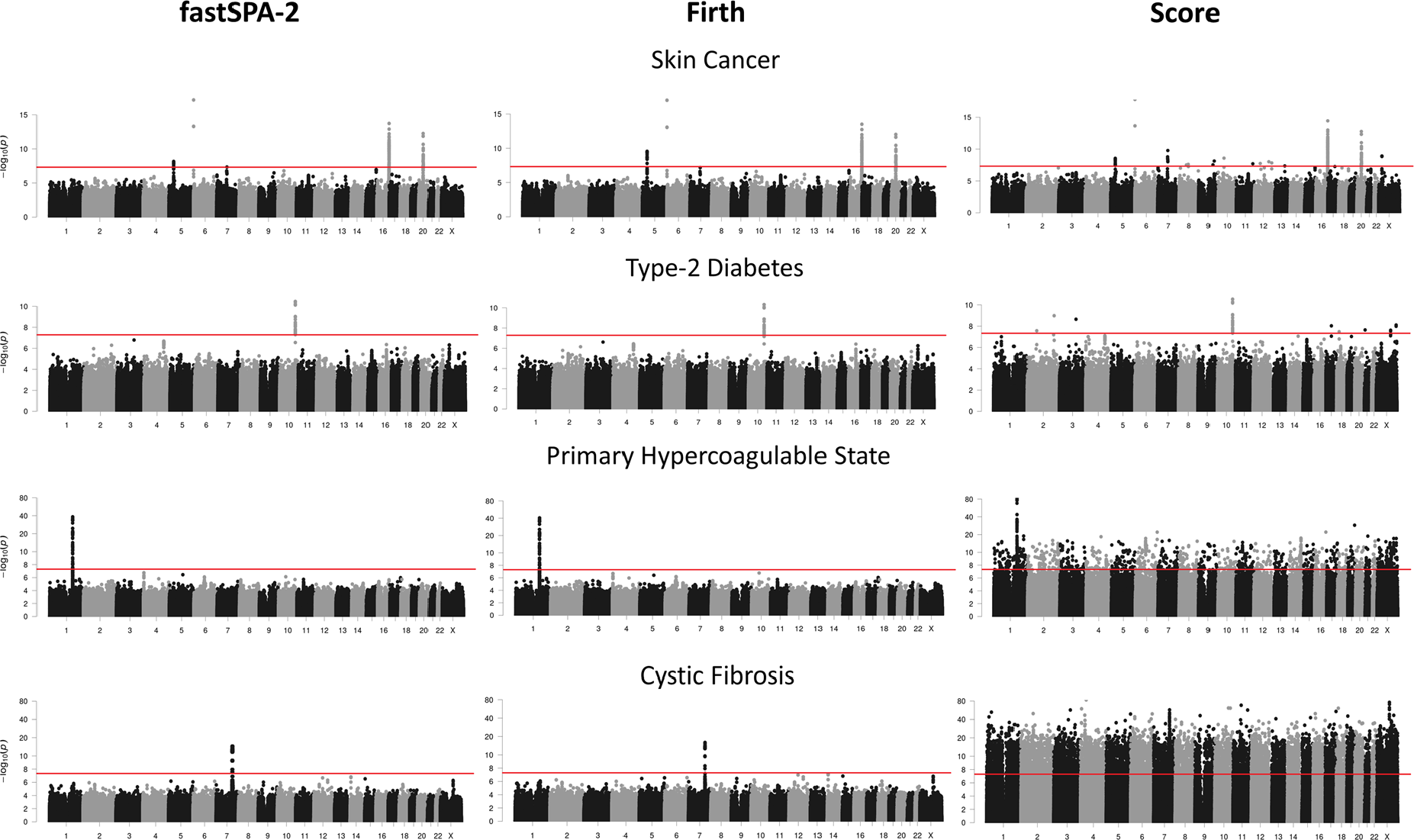
Manhattan plots for four different phenotypes from the MGI data. All genotyped variants and imputed variants with MAF > 0.001 were included in this analysis. From left to right, the three panels show associations based on the fastSPA-2, Firth, and traditional score tests, respectively. The red line represents the genome-wide significance level α = 5 × 10^−8^.

**Figure 7:**
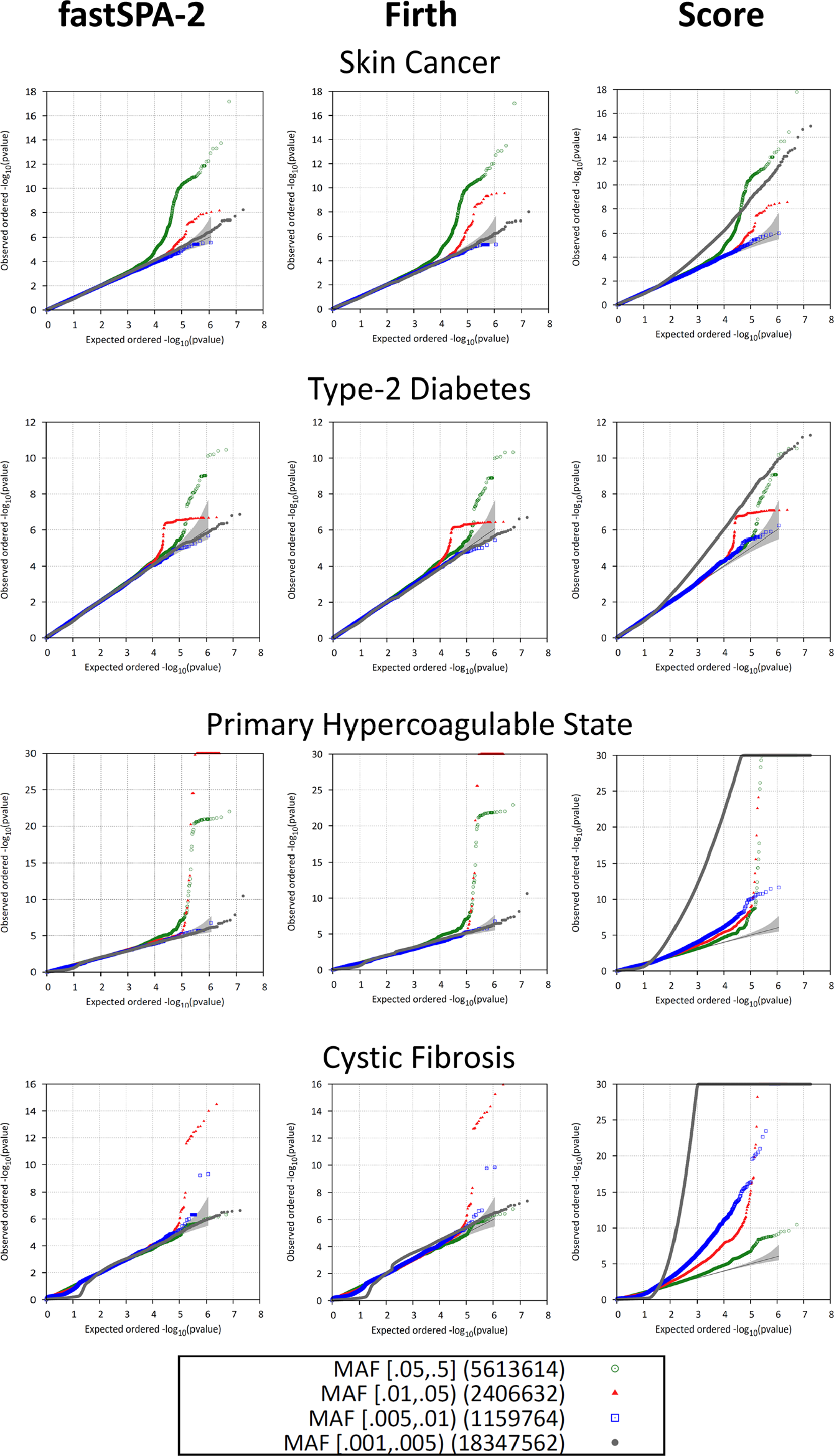
QQ plots for four different phenotypes from the MGI data. From left to right, the three panels show the QQ plots based on the fastSPA-2, Firth, and traditional score tests, respectively. The plots are color-coded based on different MAF categories.

Further, based on the *p*-values from our proposed test, we obtained the inflation factor λ of the genomic control at different *p*-value quantiles (q) and different MAF cut-offs (*Table S2*). Only the imputed variants were removed when we used different MAF cutoffs. The SNPs present on the Illumina HumanCoreExome v12.1 array were preserved. To evaluate whether using a smaller standard deviation threshold (r) improves the estimation of λ, we also applied fastSPA with r = 0.1 (i.e fastSPA-0.1), and fastSPA with the Berry-Esseen bound threshold at level *α* = 5 × 10^−8^ (fastSPA-BE) on these four phenotypes. When all the variants were included in the analysis, there was slight inflation (λ = 1.11, type 2 diabetes) or great deflation (λ = 0.12, Cystic fibrosis) at the median level for fastSPA-2. However, the genomic controls are very close to unity at q = 0.01 and q = 0.001. If we only consider the variants with MAF > 0.001, then fastSPA-2 does not show any significant inflation in λ at the median for Skin Cancer, Type-2 Diabetes, and Primary Hypercoagulable State. Although it shows a deflated genomic control for Cystic Fibrosis (λ = 0.63) due to the discrete nature of the underlying distribution. However, if we exclude the rare variants and consider only the variants with MAF > 0.01, then all four of the phenotypes show λ very close to unity. Both fastSPA-0.1 and fastSPA-BE show no significant inflation or deflation in λ at all quantiles and MAF cut-offs, except for Cystic Fibrosis (both having λ = 1.27) when all the variants are considered and genomic control is measured at the median level.

## Discussion

In this paper, we proposed a fast and scalable test to analyze large PheWAS datasets which is well calibrated even in extremely unbalanced case-control settings. The method uses computationally efficient saddle point approximation to accurately calculate p-values of score test statistics. We further proposed an improved version of our test which substantially reduces the computation time, especially for low-frequency and rare variants. Our proposed test can also adjust for additional covariates. Through extensive numerical studies we demonstrated that our test can perform 100-300 times faster than the currently used Firth’s test while retaining similar power and well controlled type I error rates. MGI data analysis illustrates that by applying the proposed method to PheWAS, we can identify true association signals while controlling for type I error, even for traits with a very small number of cases and a large number of controls.

Our test calculates *p*-values based on the traditional score test if the score statistics lie sufficiently close to the mean. Even though normal approximation is accurate near the mean, those *p*-values may not be well calibrated. In such cases, since the median *p*-values might come from the traditional score test, we can encounter slightly inflated or deflated inflation factor at median. When the case control ratio is extremely unbalanced, this phenomenon is more pronounced. One way to circumvent this issue is to measure the inflation factor at more extreme quantiles (0.01, 0.001 etc.), or to exclude rare variants when estimating the inflation factor. Another approach is to decrease the standard deviation threshold so that the median *p*-values come from the saddlepoint approximation. In the MGI data analysis, fastSPA-0.1 produced substantially improved inflation factor estimates than fastSPA-2. However, the use of threshold 0.1 instead of 2 would increase computation time ∼ 3-4x.

The Berry-Esseen threshold can be viewed as a compromise between these two thresholds. If there is no restriction in computational resource, we recommend to use fastSPA-0.1 so that most of the p-values are calculated using the saddlepoint approximation. If computational resource is limited, or researchers want to obtain results quickly, either a larger threshold (i.e fastSPA-2) or Berry-Esseen bound can be a better choice.

As sequencing cost continue to drop, whole-exome or whole-genome sequencing will be used for PheWAS to identify rare variants associated with clinical phenotypes.^31^ In rare variant association analysis, gene or region based multiple variant tests are commonly used to improve power.^32^ When case-control ratios are unbalanced, popular rare variant tests, including burden tests, SKAT and SKAT-O, can also have substantially inflated type I error rates. Although resampling based approaches have been developed to address this problem,^33^ the existing methods are not fast enough to be used in PheWAS. One possible approach is first to adjust single variant score statistics using SPA and then to use the adjusted score statistics to control for the type I error. We left it for future research.

In summary, we have proposed an accurate and scalable method for PheWAS data analysis. With the growing effort to build large research cohorts for precision medicine^31^, future PheWAS would have hundreds of thousands of samples and hundreds of millions of variants. Our method will provide a scalable solution for this large-scale problem and contribute to finding genetic component of complex traits. All our tests are implemented in the R package **SPAtest.**

## Supplemental Data

Four additional figures and two additional tables that were referred to in this paper, are provided in the supplemental data.

## Acknowledgements

This work was supported by grants R01 HG008773 (RD and SL). We would like to thank the investigators of MGI project for access to the PheWAS dataset and Dr. Hyun Min Kang for implementing the methods in the Epacts package.

## Web resources

SPAtest R-package: https://sites.google.com/a/umich.edu/leeshawn/software

Michigan Genomics Initiative:https://www.michigangenomics.org/

Online Mendelian Inheritance in Man (OMIM): http://www.omim.org

## Appendix

### Explanation behind using 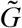 instead of *G*

We first note that 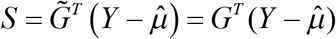 since 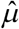 is the maximum likelihood estimator of *μ* under the null model and 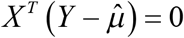. Now, the score function and the observed information matrix under the null model are given by,

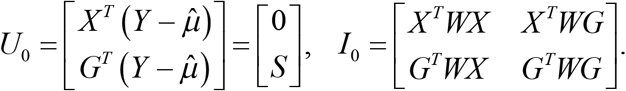

Therefore, the variance of *S* under *H*_0_ is given by,

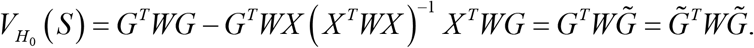

So, even though the two expressions of *S* are algebraically the same, the variance can be expressed as a weighted sum of 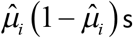 where the weights are given by 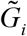 s. Therefore, we used 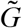 instead of *G* to express the score statistic.

## References

1. Welter, D., MacArthur, J., Morales, J., Burdett, T., Hall, P., Junkins, H., Klemm, A., Flicek, P., Manolio, T., Hindorff, L., et al., (2014). The NHGRI GWAS Catalog, a curated resource of SNP-trait associations. Nucleic Acids Res 42, D1001–1006.

2. Solovieff, N., Cotsapas, C., Lee, P.H., Purcell, S.M., and Smoller, J.W. (2013). Pleiotropy in complex traits: challenges and strategies. Nat Rev Genet 14, 483–495.

3. Denny, J.C., Ritchie, M.D., Basford, M.A., Pulley, J.M., Bastarache, L., Brown-Gentry, K., Wang, D., Masys, D.R., Roden, D.M., and Crawford, D.C. (2010). PheWAS: demonstrating the feasibility of a phenome-wide scan to discover gene-disease associations. Bioinformatics 26, 1205–1210.

4. Denny, J.C., Crawford, D.C., Ritchie, M.D., Bielinski, S.J., Basford, M.A., Bradford, Y., Chai, H.S., Bastarache, L., Zuvich, R., Peissig, P., et al., (2011). Variants near FOXE1 are associated with hypothyroidism and other thyroid conditions: using electronic medical records for genome-and phenome-wide studies. Am J Hum Genet 89, 529–542.

5. Hebbring, S.J., Schrodi, S.J., Ye, Z., Zhou, Z., Page, D., and Brilliant, M.H. (2013). A PheWAS approach in studying HLA-DRB1*1501. Genes Immun 14, 187–191.

6. Ritchie, M.D., Denny, J.C., Zuvich, R.L., Crawford, D.C., Schildcrout, J.S., Bastarache, L., Ramirez, A.H., Mosley, J.D., Pulley, J.M., Basford, M.A., et al., (2013). Genome- and phenome-wide analyses of cardiac conduction identifies markers of arrhythmia risk. Circulation 127, 1377–1385.

7. Pendergrass, S.A., Brown-Gentry, K., Dudek, S., Frase, A., Torstenson, E.S., Goodloe, R., Ambite, J.L., Avery, C.L., Buyske, S., Buzkova, P., et al., (2013). Phenome-wide association study (PheWAS) for detection of pleiotropy within the Population Architecture using Genomics and Epidemiology (PAGE) Network. PLoS Genet 9, e1003087.

8. Shameer, K., Denny, J.C., Ding, K., Jouni, H., Crosslin, D.R., de Andrade, M., Chute, C.G., Peissig, P., Pacheco, J.A., Li, R., et al., (2014). A genome- and phenome-wide association study to identify genetic variants influencing platelet count and volume and their pleiotropic effects. Hum Genet 133, 95–109.

9. Hebbring, S.J. (2014). The challenges, advantages and future of phenome-wide association studies. Immunology 141, 157–165.

10. Marchini, J., and Howie, B. (2010). Genotype imputation for genome-wide association studies. Nat Rev Genet 11, 499–511.

11. Cox, D., and Hinkley, D. (1974). Theoretical Statistics.(London: Chapman and Hall).

12. Ma, C., Blackwell, T., Boehnke, M., Scott, L.J., and the Go, T.D.i. (2013). Recommended Joint and Meta-Analysis Strategies for Case-Control Association Testing of Single Low-Count Variants. Genetic Epidemiology 37, 539–550.

13. Firth, D. (1993). Bias reduction of maximum likelihood estimates. Biometrika 80, 27–38.

14. Daniels, H.E. (1954). Saddlepoint Approximations in Statistics. Ann Math Stat 25, 631–650.

15. Barndorff-Nielsen, O.E. (1990). Approximate Interval Probabilities. Journal of the Royal Statistical Society Series B (Methodological) 52, 485–496.

16. Kuonen, D. (1999). Saddlepoint approximations for distributions of quadratic forms in normal variables. Biometrika 86, 929–935.

17. Feller, W. (1945). The fundamental limit theorems in probability. 800–832.

18. Berry, A.C. (1941). The accuracy of the Gaussian approximation to the sum of independent variates. T Am Math Soc 49, 122–136.

19. Esseen, C.G. (1942). On the Liapounoff Limit of Error in the Theory of Probability. Ark Mat Astr och Fys 28A, 1–19.

20. Esseen, C.G. (1956). A Moment Inequality with an Application to the Central Limit Theorem. Skand Aktuarietidskr 39, 160–170.

21. Jensen, J.L. (1995). Saddlepoint Approximations.(Oxford: Oxford University Press).

22. Whittaker, E.T., and Robinson, G. (1967). The Newton-Raphson Method. In The Calculus of Observations: A Treatise on Numerical Mathematics. (New York, Dover), pp 84–87.

23. Press, W.H.F., B. P.; Teukolsky, S. A.; and Vetterling, W. T. (1992). Numerical Recipes in FORTRAN: The Art of Scientific Computing.(Cambridge, England: Cambridge University Press).

24. Brent, R.P. (1973). Algorithms for Minimization without Derivatives.(Englewood Cliffs, NJ: Prentice-Hall).

25. Shevtsova, I.G. (2010). An improvement of convergence rate estimates in the Lyapunov theorem. Doklady Mathematics 82, 862–864.

26. McCarthy, S., Das, S., Kretzschmar, W., Delaneau, O., Wood, A.R., Teumer, A., Kang, H.M., Fuchsberger, C., Danecek, P., Sharp, K., et al., (2016). A reference panel of 64,976 haplotypes for genotype imputation. Nat Genet 48, 1279–1283.

27. Delaneau, O., Zagury, J.-F., and Marchini, J. (2013). Improved whole-chromosome phasing for disease and population genetic studies. Nat Meth 10, 5–6.

28. Loh, P.-R., Danecek, P., Palamara, P.F., Fuchsberger, C., A Reshef, Y., K Finucane, H., Schoenherr, S., Forer, L., McCarthy, S., Abecasis, G.R., et al., (2016). Reference-based phasing using the Haplotype Reference Consortium panel. Nat Genet 48, 1443–1448.

29. Das, S., Forer, L., Schonherr, S., Sidore, C., Locke, A.E., Kwong, A., Vrieze, S.I., Chew, E.Y., Levy, S., McGue, M., et al., (2016). Next-generation genotype imputation service and methods. Nat Genet 48, 1284–1287.

30. Carroll, R.J., Bastarache, L., and Denny, J.C. (2014). R PheWAS: data analysis and plotting tools for phenome-wide association studies in the R environment. Bioinformatics 30, 2375–2376.

31. Collins, F.S., and Varmus, H. (2015). A New Initiative on Precision Medicine. New England Journal of Medicine 372, 793–795.

32. Lee, S., Abecasis, Gonçalo R., Boehnke, M., and Lin, X. (2014). Rare-Variant Association Analysis: Study Designs and Statistical Tests. American Journal of Human Genetics 95, 5–23.

33. Lee, S., Fuchsberger, C., Kim, S., and Scott, L. (2015). An efficient resampling method for calibrating single and gene-based rare variant association analysis in case-control studies. Biostatistics 17, 1–15.

34. Zhang, M., Song, F., Liang, L., Nan, H., Zhang, J., Liu, H., Wang, L.-E., Wei, Q., Lee, J.E., Amos, C.I., et al., (2013). Genome-wide association studies identify several new loci associated with pigmentation traits and skin cancer risk in European Americans. Hum Mol Genet 22, 2948–2959.

35. Sulem, P., Gudbjartsson, D.F., Stacey, S.N., Helgason, A., Rafnar, T., Magnusson, K.P., Manolescu, A., Karason, A., Palsson, A., Thorleifsson, G., et al., (2007). Genetic determinants of hair, eye and skin pigmentation in Europeans. Nat Genet 39, 1443–1452.

36. Jacobs, L.C., Hamer, M.A., Gunn, D.A., Deelen, J., Lall, J.S., van Heemst, D., Uh, H.-W., Hofman, A., Uitterlinden, A.G., Griffiths, C.E.M., et al., (2015). A Genome-Wide Association Study Identifies the Skin Color Genes IRF4, MC1R, ASIP, and BNC2 Influencing Facial Pigmented Spots. J Invest Dermatol 135, 1735–1742.

37. Liu, F., Visser, M., Duffy, D.L., Hysi, P.G., Jacobs, L.C., Lao, O., Zhong, K., Walsh, S., Chaitanya, L., Wollstein, A., et al., (2015). Genetics of skin color variation in Europeans: genome-wide association studies with functional follow-up. Human genetics 134, 823–835.

38. Barrett, J.H., Iles, M.M., Harland, M., Taylor, J.C., Aitken, J.F., Andresen, P.A., Akslen, L.A., Armstrong, B.K., Avril, M.-F., Azizi, E., et al., (2011). Genome-wide association study identifies three new melanoma susceptibility loci. In Nat Genet. pp 1108–1113.

39. Nan, H., Kraft, P., Qureshi, A.A., Guo, Q., Chen, C., Hankinson, S.E., Hu, F.B., Thomas, G., Hoover, R.N., Chanock, S., et al., (2009). Genome-wide association study of tanning phenotype in a population of European ancestry. J Invest Dermatol 129, 2250–2257.

40. Scott, L.J., Bonnycastle, L.L., Willer, C.J., Sprau, A.G., Jackson, A.U., Narisu, N., Duren, W.L., Chines, P.S., Stringham, H.M., Erdos, M.R., et al., (2006). Association of transcription factor 7-like 2 (TCF7L2) variants with type 2 diabetes in a Finnish sample. Diabetes 55, 2649–2653.

41. Bertina, R.M., Koeleman, B.P., Koster, T., Rosendaal, F.R., Dirven, R.J., de Ronde, H., van der Velden, P.A., and Reitsma, P.H. (1994). Mutation in blood coagulation factor V associated with resistance to activated protein C. Nature 369, 64–67.

42. Kerem, B., Rommens, J.M., Buchanan, J.A., Markiewicz, D., Cox, T.K., Chakravarti, A., Buchwald, M., and Tsui, L.C. (1989). Identification of the cystic fibrosis gene: genetic analysis. Science 245, 1073–1080.

